# Diminished spatial dynamics and maladaptive spatial complexity link resting brain network disruption to cognition in schizophrenia

**DOI:** 10.64898/2025.12.07.692856

**Authors:** Krishna Pusuluri, Aysenil Belger Arcasoy, Armin Iraji, Vince D. Calhoun

## Abstract

Resting-state fMRI studies increasingly emphasize the dynamic nature of brain networks. While most approaches examine temporal fluctuations in connectivity, we focus on the spatial dynamics and complexity at voxel level - how networks expand and contract, and change their structural complexity over time. Using dynamic independent component analysis (ICA), we investigate the hierarchical structure of the resulting time-varying spatial networks, from their broad periphery to their most active core. We combine this with fractal dimension (FrD) as a measure of a network’s spatial complexity and analyze temporal changes (dynamic flexibility) in a network and synchronized fluctuations between network pairs (fractal dimension coupling, FrDC). We refer to this approach as “dynamic spatial network complexity and connectivity (dSNCC)”. Using a combined cohort of 508 subjects (315 healthy controls, 193 schizophrenia patients), we found that schizophrenia is associated with higher mean FrD in several networks, suggesting more irregular patterns/boundaries and a disorganized network structure. Critically, patients showed significantly reduced dynamic flexibility, indicating their networks are “stuck” in a less adaptable state. This robust finding is evidenced by a synergistic loss of temporal standard deviations in both network volume and FrD across multiple networks and activity thresholds. This maladaptive complexity was associated with cognitive impairment, with several dSNCC measures showing significant associations with subject scores for processing speed, visual learning, and verbal learning. Higher complexity in these networks and more significantly, their reduced dynamic flexibility as seen in patients, were particularly associated with impaired performance. Furthermore, we found aberrant connectivity (FrDC) in schizophrenia, with certain network pairs exhibiting overly synchronized complexity changes. Our results demonstrate that dSNCC is a powerful tool for characterizing network dynamics and may potentially provide a measurable mechanism for maladaptation in schizophrenia, where the brain’s inability to fluidly change its complexity may contribute to cognitive deficits and symptoms like disorganized thought. These findings highlight the importance of studying the intrinsic spatial dynamic properties to reveal the fundamental principles of brain network organization in health and disease. Our work represents a significant leap in complex systems neuroscience and provides a novel, quantifiable biomarker framework highly relevant for understanding and targeting other complex disorders characterized by network dysfunction, such as Alzheimer’s disease, autism, or other mental health conditions.

## 1 Introduction

The field of neuroimaging has increasingly shifted toward a dynamic view of brain function, moving away from the traditional focus on static brain networks to explore how network relationships change over time [1, 2, 3]. Functional magnetic resonance imaging (fMRI), and particularly resting-state fMRI (rsfMRI), has been a pivotal tool in this evolution, allowing researchers to investigate the brain’s intrinsic activity and identify large-scale networks with coherent spatiotemporal activity patterns [4]. While a common assumption in many studies is that the spatial patterns of these networks remain fixed, this overlooks the inherent temporal evolution of brain networks at voxel level, where they can expand, shrink, or change in their spatial complexity.

It was shown that schizophrenia (SZ) is strongly associated with the temporal dynamics of whole-brain network connectivity, revealing transient states of dysconnectivity [4]. The current study extends this focus to the spatial dynamics of these networks, arguing that the true complexity of brain function lies not only in how networks connect but also in how they change their intrinsic spatial organization over time [5, 6]. We previously developed methods to quantify these spatial dynamics, demonstrating that features like network volume and volumetric coupling (synchronized growth and shrinkage) are altered in SZ and associated with cognitive performance [7, 8]. Building upon this foundation, we now extend our analysis to investigate the complexity of these dynamic spatial brain networks. Fractal geometry provides a powerful tool for characterizing the intricate, self-similar patterns often found in biological systems, offering a measure of complexity that goes beyond simple volume. The brain’s architecture, from the gyri and sulci to the branching of neurons, exhibits fractal properties, suggesting that fractal analysis is a particularly relevant approach for studying brain networks [9, 10, 11, 12]. Complexity has been successfully used to study cortical structure in various brain disorders, including SZ [9, 13, 14, 15, 16, 17]. For instance, previous studies have shown that changes in cortical complexity, as measured by fractal dimension (FrD), are linked to age-related cognitive decline and neurodevelopmental disorders. By investigating the complexity of these networks dynamically, we aim to provide a quantitative measure which could serve as a novel and powerful biomarker for SZ and other brain disorders. We also extend our previous analysis on volumetric changes of these networks and their group differences by including more covariates in the analysis to increase model accuracy and account for confounding effects [7]. We then seek to link these fundamental network properties to cognitive performance, ultimately helping to inform more personalized and effective treatment strategies.

We employ dynamic independent component analysis (ICA) [18] to evaluate spatially dynamic brain networks and probe the hierarchical organization of brain activity - from broad, low-activity regions to focused, highly active cores - providing a more nuanced view of network dynamics. We use a box-counting method for FrD to quantify the temporal changes in network complexity [14, 17]. We refer to this approach of combining windowed ICA with FrD to measure complexity and coupling as dynamic spatial network complexity and connectivity (dSNCC) and evaluate how these are associated with SZ and cognitive function. We hypothesize that the fractal complexity of brain networks and the coupling of this complexity between networks will reveal novel insights into the neuropathology of SZ. Specifically, we expect that SZ will be associated with altered FrD and fractal dimension coupling (FrDC), both of which will be linked to cognitive impairment.

## 2 Methods

### 2.1 Data collection and preprocessing

We used 3-Tesla resting-state functional magnetic resonance imaging (rsfMRI) data from a combined cohort of 508 subjects (315 controls and 193 SZ patients) from the FBIRN (Functional Imaging Biomedical Informatics Research Network) [4], COBRE (Center for Biomedical Research Excellence) [19], and MPRC (Maryland Psychiatric Research Center) [20] datasets. Preprocessing of rsfMRI data was performed using the statistical parametric mapping toolbox (SPM12). This included standard procedures for motion correction, slice-timing correction, and spatial normalization to the Montreal Neurological Institute template. Further details on the datasets and preprocessing can be found in our previous work [7, 18].

### 2.2 Analysis pipeline

Our analysis pipeline, depicted in Figure 1, is adapted from our previous volumetric study to incorporate complexity analysis. In the first step, group-level spatially constrained independent component analysis (sICA) was performed on the rsfMRI data with a model order of 20 components using the GIFT software package (https://trendscenter.org/software/gift/) [3]. This process identified 14 relevant intrinsic connectivity networks (ICNs) as shown in Figure 2 and their associated time courses. In the second step, a sliding-window approach combined with spatially constrained ICA (MOOICAR - multiobjective optimization ICA with reference) was performed for each subject with a sliding window length of 60s [7, 18] to capture the temporal evolution of the spatial patterns of these networks while maintaining the correspondence of ICNs across subjects and time windows.

**Figure 1:**
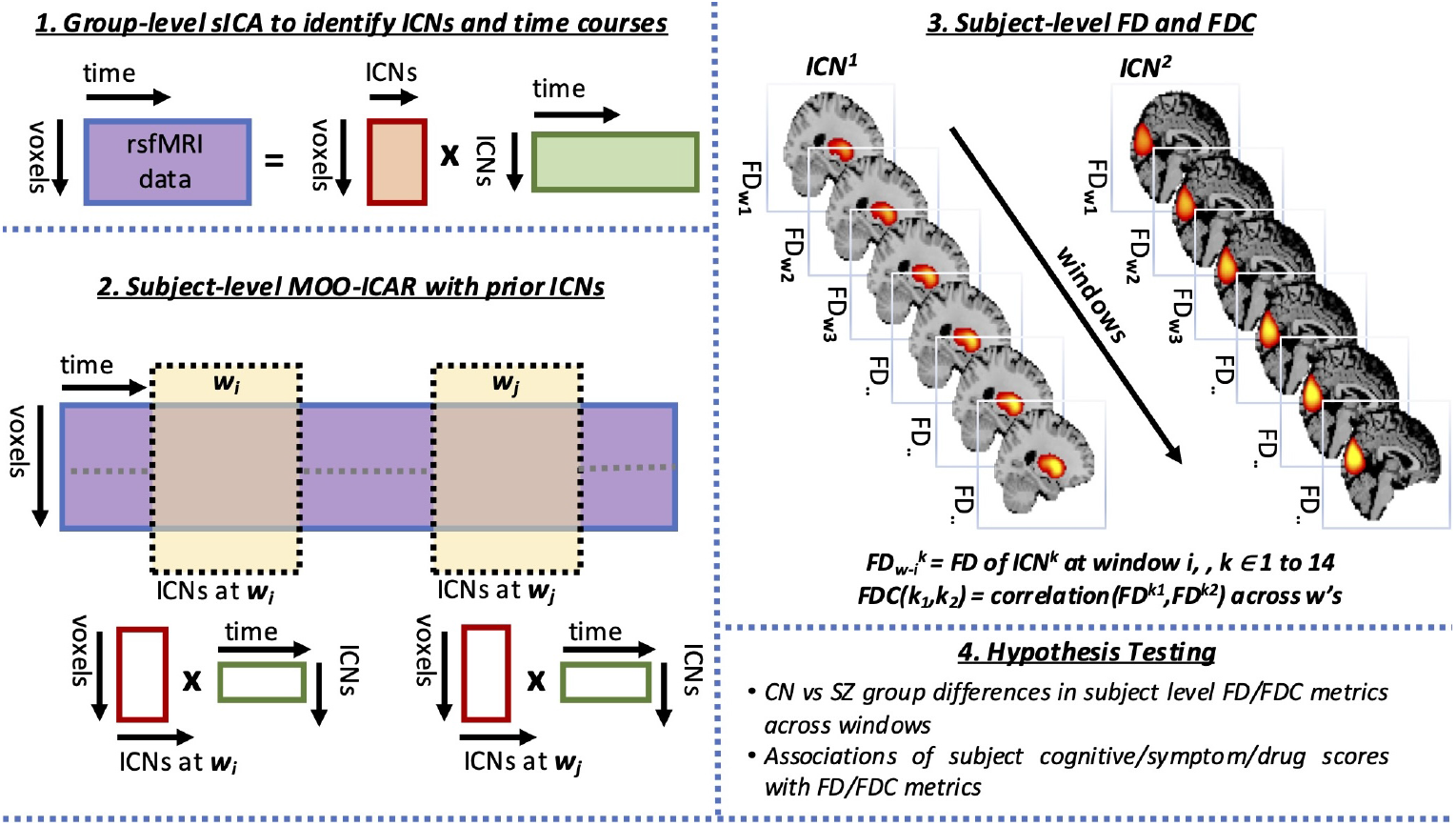
Analysis Pipeline. A diagram showing the analysis pipeline: (1) group-level sICA is performed to identify intrinsic connectivity networks (ICNs) and their time courses; (2) subject-level MOOICAR is performed with prior ICNs to get spatially dynamic ICNs for each subject across sliding windows; (3) the fractal dimension (FrD) for each ICN and fractal dimension coupling (FrDC) between network pairs are computed for each subject across windows at different activity thresholds (*Z*_*th*_) and (4) hypothesis testing is performed to study group differences and associations with subject scores.

**Figure 2:**
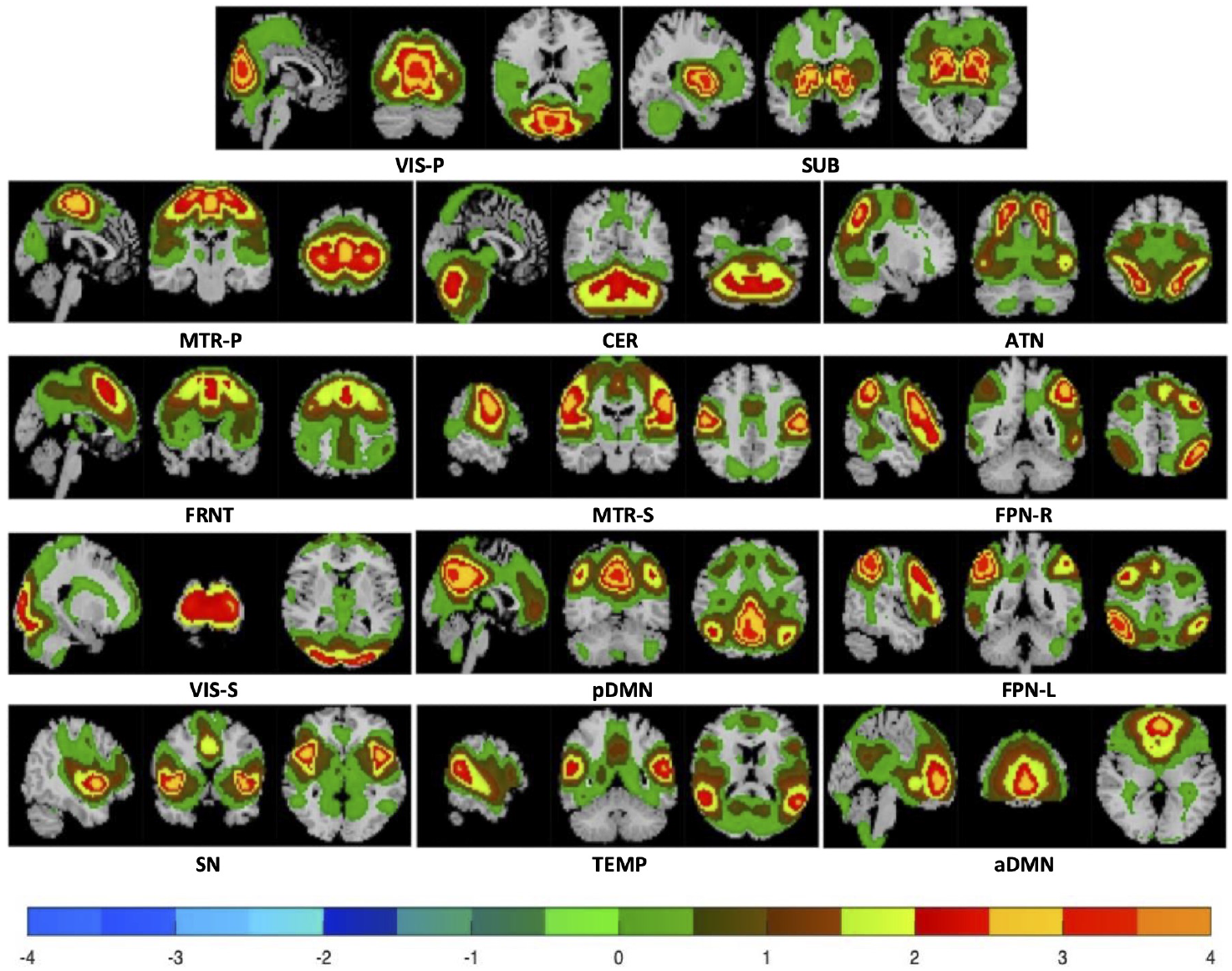
Hierarchical network activity maps. Mean activity maps (CN group) for each of the 14 relevant brain networks are shown at different activity thresholds. The z-scored maps use a discrete colormap to illustrate brain network activity, with a threshold of *Z*_*th*_=0 showing broad regions of above-average activity and higher thresholds in steps of 0.5 unveiling progressively more focused, highly active core regions. The images are displayed on three planar cross sections—sagittal, coronal, and transverse. The networks shown are: VIS-P (visual primary), SUB (subcortical), MTR-P (somatomotor primary), CER (cerebellar), ATN (attention-dorsal), FRNT (frontal), MTR-S (somatomotor secondary), FPN-R (frontoparietal right), VIS-S (visual secondary), pDMN (posterior default mode), FPN-L (frontoparietal left), SN (salience), TEMP (temporal), and aDMN (anterior default mode).

In the third step, for each subject and time window, we calculated the FrD of each ICN using a box-counting method as shown in Figure 3. This method involves covering the network’s active voxels with a grid of boxes of varying sizes and counting the number of boxes required to cover the network (Figure 3 A-D). The FrD is then estimated from the slope of a log-log plot of the number of boxes versus the box size (Figure 3E). We performed this analysis at various activity thresholds (*Z*_*th*_ = 0, 0.5, 1, 1.5, 2, 2.5, 3, 3.5) such that for a chosen value of the threshold *Z*_*th*_, all the voxels with z-scored activity above this threshold are considered. This allowed us to examine the hierarchical organization of complexity starting from the wide-spread regions of above average activity (*Z*_*th*_=0) to the most active, focused regions of each network at higher thresholds. The temporal variations in FrD across windows for a subject at a given threshold are computed as shown in Figure 3F, from which we computed the subject-level mean and standard deviations (std) of FrD. We also computed the subject-level FrDC between network pairs by correlating their FrDs across time windows (correlations are converted to Fisher’s z-scores). This yields a 14 × 14 FrDC matrix for each subject at each threshold, where a large positive coupling suggests that two networks grow and shrink in complexity together, while a negative value implies a complementary relationship.

**Figure 3:**
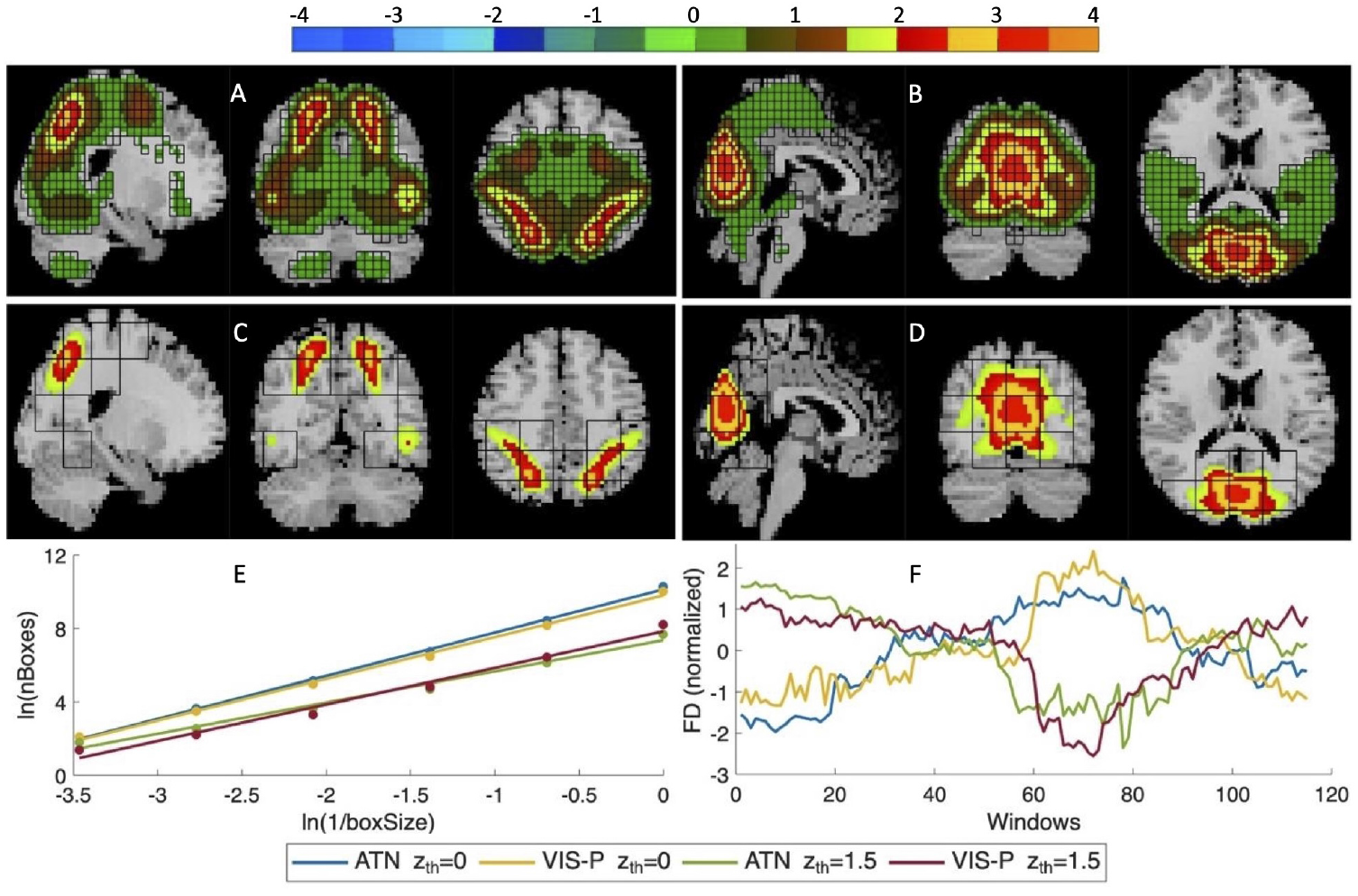
Demonstration of fractal dimension (FrD) via box counting and temporal changes in FrD. This figure demonstrates the process of calculating FrD using the box-counting method. Panels A-D show mean CN group network maps (across subjects/windows) for the following combinations, with a box grid overlaid to illustrate the counting process: A (ATN network at *Z*_*th*_=0 with a box size of 2), B (VIS-P network at *Z*_*th*_=0 with a box size of 2), C (ATN network at *Z*_*th*_=1.5 with a box size of 8), and D (VIS-P network at *Z*_*th*_=1.5 with a box size of 8). Box counting (number of boxes with active voxels) is repeated at different box sizes to compute FrD at a given threshold, as shown in panel E with the log-log plot of the number of active boxes versus the inverse of the box size (ln represents natural logarithm). The slopes of the regression lines in panel E represent the FrD for the mean network maps shown in A (FrD=2.35), B (FrD=2.28), C (FrD=1.7), and D (FrD=1.99). Panel F illustrates the temporal changes in FrD for these network and threshold combinations, normalized across time windows for a single subject.

Finally, in the fourth step, we used the subject-level FrD and FrDC matrices to perform hypothesis testing. We used robust linear regressions with age, sex, and site as covariates (as shown in Figures 4 5, 6, 7, 8C, 8F) to find group differences in subject-level mean and standard deviation (across windows) of volumes, FrDs, and FrDC metrics between SZ patients and healthy controls (CN). A 5% false discovery rate (FDR) was used to correct for multiple comparisons [21], with FDR correction applied across all thresholds or each threshold separately. We also performed robust linear regression analysis to investigate the association between subject-level FrD and FrDC metrics with cognitive (visual learning, verbal learning, processing speed, working memory), symptom (positive and negative syndrome scale - PANSS), and drug scores (Chlorpromazine - CPZ - equivalent). Further details of these scores and associations can be found in [7].

**Figure 4:**
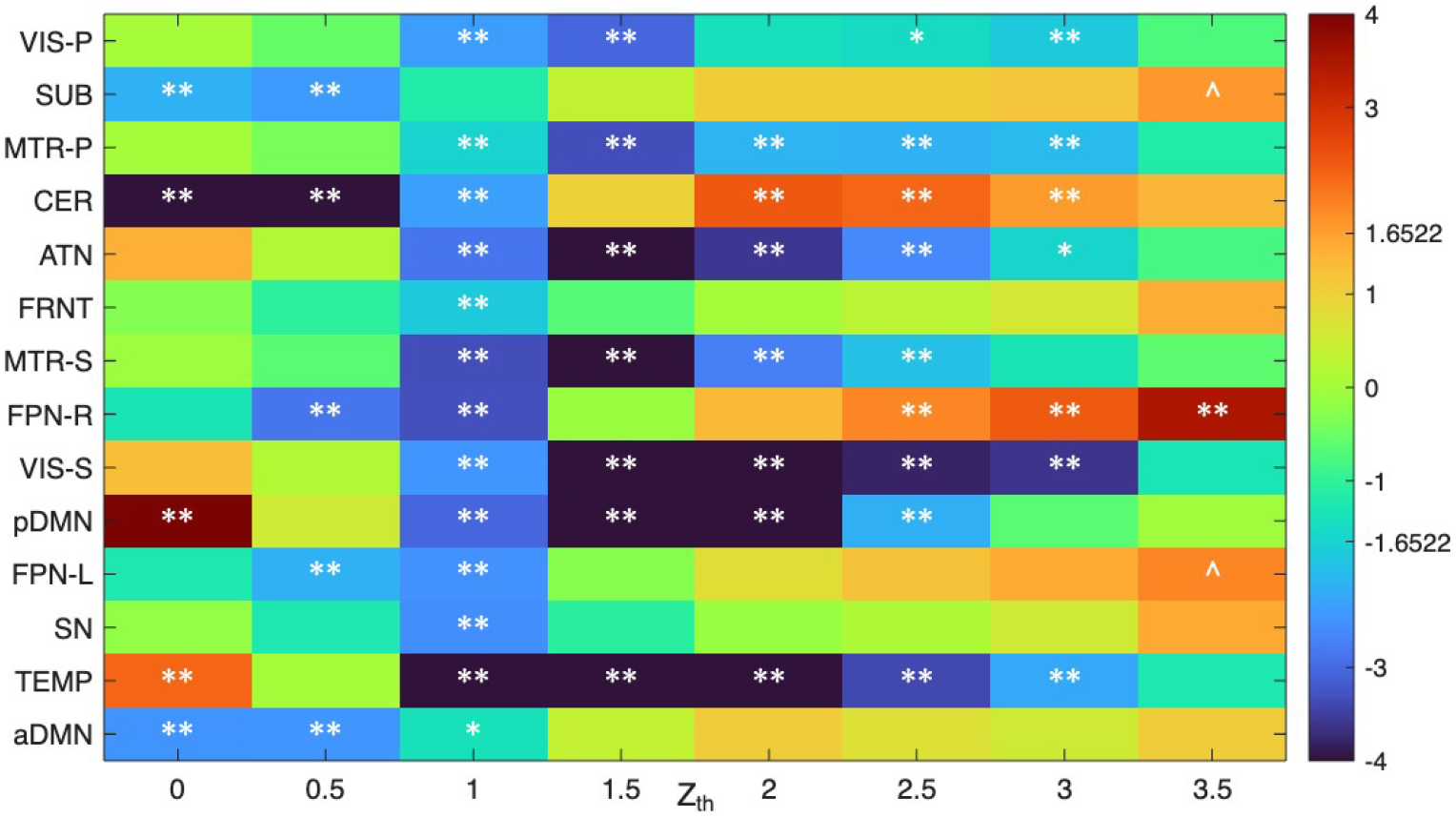
Group differences (CN-SZ) in subject level mean fractal dimension (FrD) of networks across windows. A heatmap depicting robust regression results with age, sex and site as covariates for group differences in subject-level mean FrD between the control (CN) and schizophrenia (SZ) groups across different activity thresholds (*Z*_*th*_). The color bar represents the t-statistic values (k=-log10(p) × sign(t)), where t is the ratio of the regression coefficient to the standard error of its estimate. Here positive values (red) indicate networks with higher mean FrD in CN than SZ, and negative values (blue) indicate networks with higher mean FrD in SZ than CN. Significant results are denoted by symbols: ∗∗ (significant with FDR correction at both single threshold as well as across all thresholds), ∗ (significant with FDR at a single threshold), and ^(significant with FDR at all thresholds but not a single threshold) (^-while generally less likely, this remains statistically possible contingent upon the specific set of p-values involved in the FDR calculation across individual or combined thresholds). Overall, the mean FrD for significant results is predominantly higher in SZ than CN. Critical value (1.6522) for significance across thresholds is shown on the colormap.

**Figure 5:**
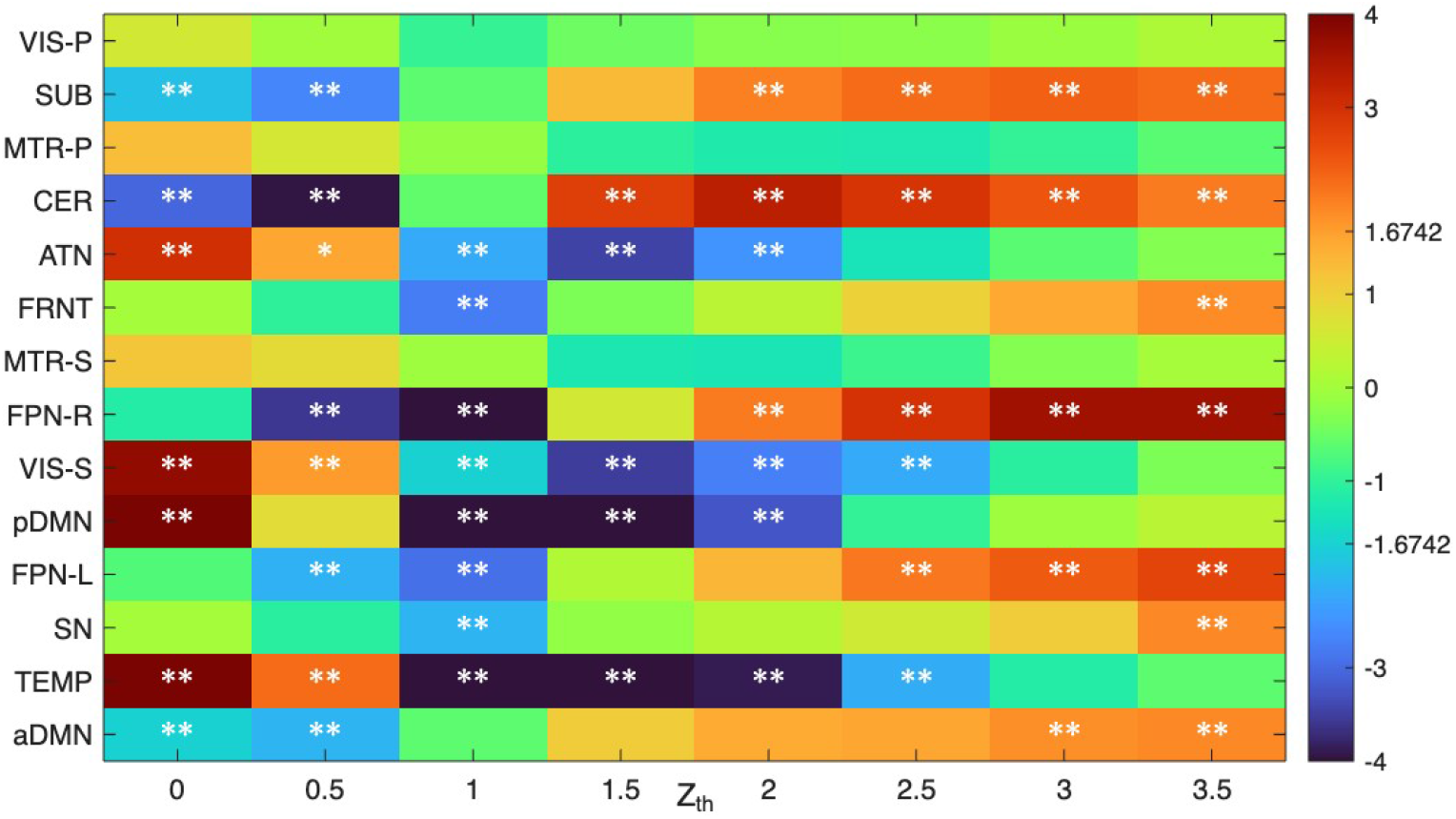
Group differences (CN-SZ) in subject level mean volume of networks across windows. A heatmap depicting robust regression results with age, sex and site as covariates for group differences in subject-level mean volume between the control (CN) and schizophrenia (SZ) groups across different activity thresholds (*Z*_*th*_). The color bar represents the t-statistic values (k=-log10(p) × sign(t)), where positive values (red) indicate networks with higher mean volume in CN than SZ, and negative values (blue) indicate networks with higher mean volume in SZ than CN. Significant results are denoted by symbols: ∗∗ (significant with FDR correction at both single threshold as well as across all thresholds), ∗ (significant with FDR at a single threshold), and ^(significant with FDR at all thresholds but not a single threshold). Critical value (1.6742) for significance across thresholds is shown on the colormap.

**Figure 6:**
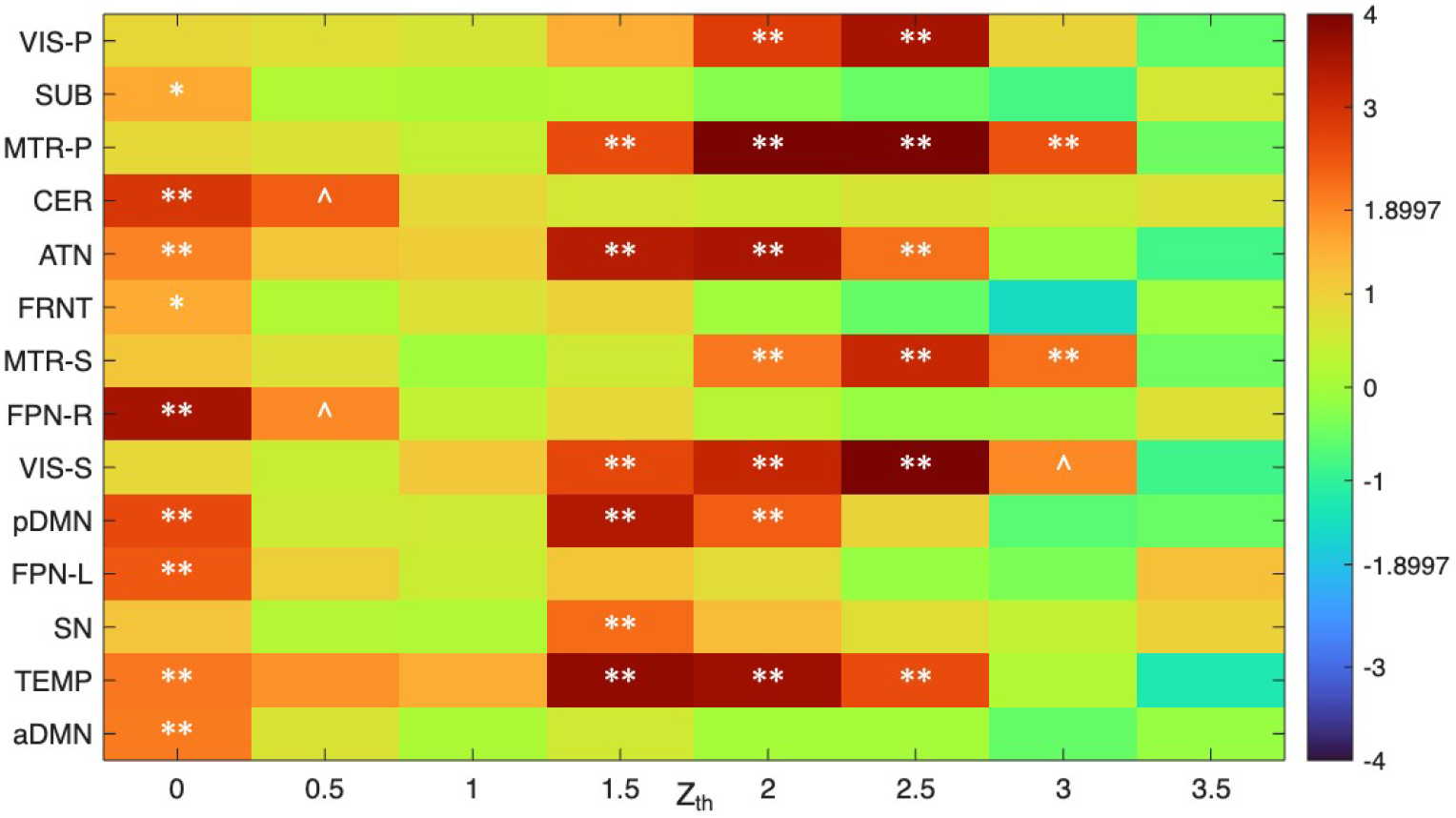
Group differences (CN-SZ) in subject level standard deviation (std.) of fractal dimension (FrD) of networks across windows. A heatmap depicting robust regression results with age, sex and site as covariates for group differences in the subject-level std. of FrD across windows between the control (CN) and schizophrenia (SZ) groups across different activity thresholds (*Z*_*th*_). The color bar represents the t-statistic values (k=-log10(p) × sign(t)), where positive values (red) indicate networks with higher std. of FrD in CN than SZ, and negative values (blue) indicate networks with higher std. of FrD in SZ than CN. Significant results are denoted by symbols: ** (significant with FDR correction at both a single threshold as well as across all thresholds), * (significant with FDR at a single threshold), and ^(significant with FDR at all thresholds but not a single threshold). The results consistently show that std. of FrD is less in SZ than in CN, suggesting reduced flexibility in network dynamics. Critical value (1.8997) for significance across thresholds is shown on the colormap.

**Figure 7:**
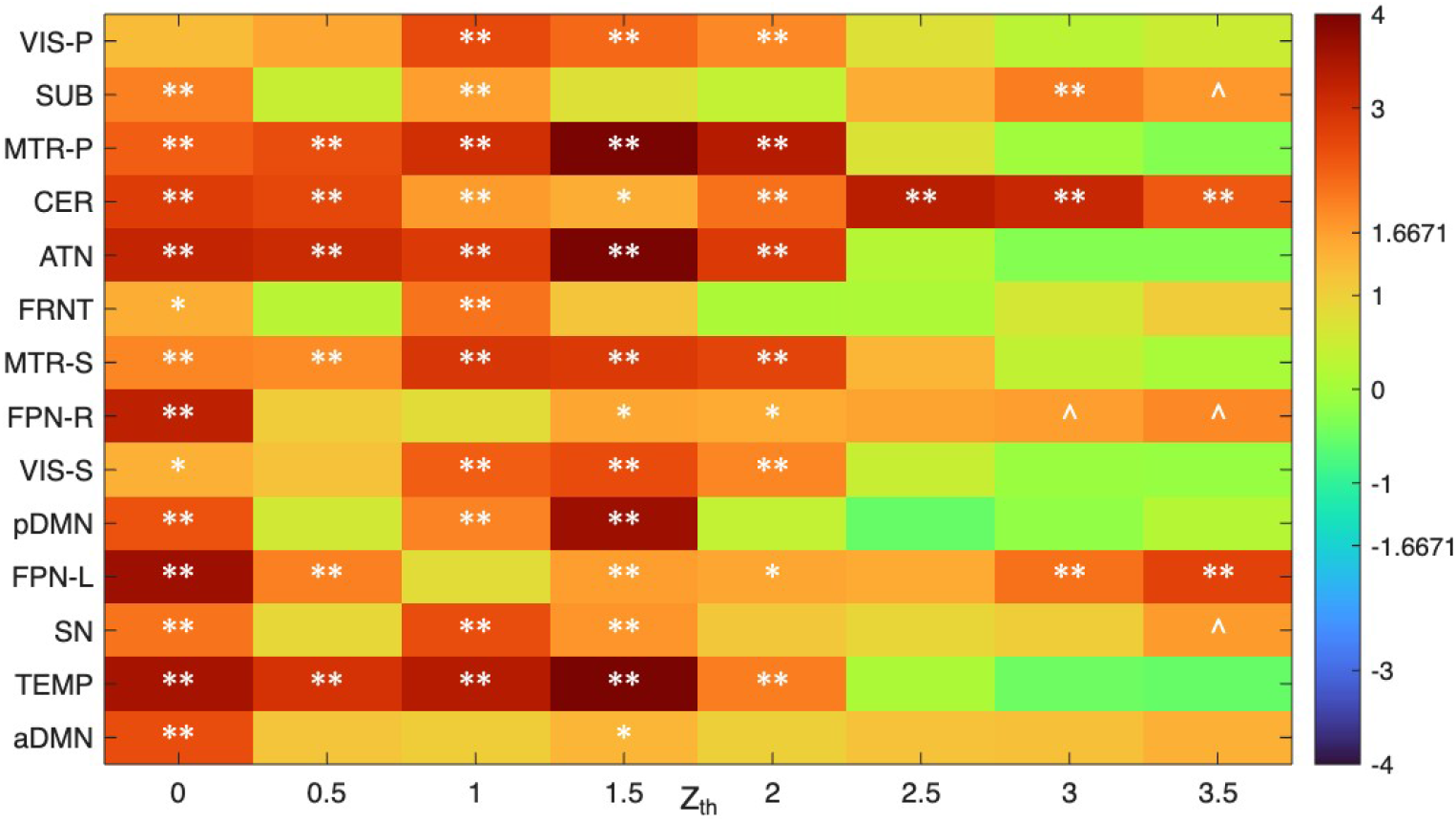
Group differences (CN-SZ) in subject level standard deviation (std.) of volume of networks across windows. A heatmap depicting robust regression results with age, sex and site as covariates for group differences in the subject-level std. of volume across windows between the control (CN) and schizophrenia (SZ) groups across different activity thresholds (*Z*_*th*_). The color bar represents the t-statistic values (k=-log10(p) × sign(t)), where positive values (red) indicate networks with higher std. of volume in CN than SZ, and negative values (blue) indicate networks with higher std. of FrD in SZ than CN. Significant results are denoted by symbols: ** (significant with FDR correction at both a single threshold as well as across all thresholds), * (significant with FDR at a single threshold), and ^(significant with FDR at all thresholds but not a single threshold). The results consistently show that std. of volume is less in SZ than in CN, suggesting reduced flexibility in network dynamics. Critical value (1.6671) for significance across thresholds is shown on the colormap.

**Figure 8:**
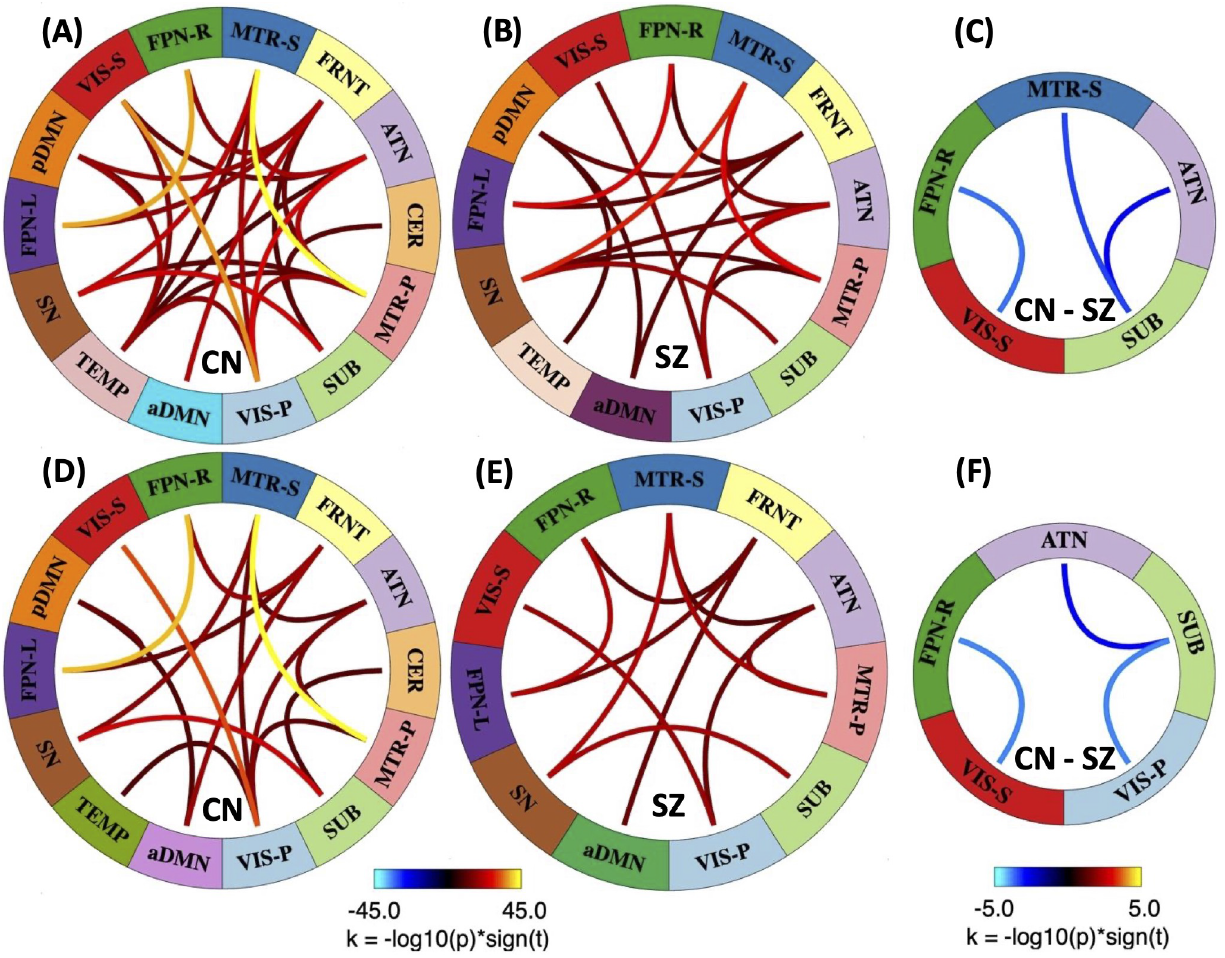
Connectograms showing 1-sample results for significant fractal dimension coupling (FrDC) and (CN-SZ) group differences at Z_th_ = 0 and Z_th_ = 0.5. Panels A, B, and C show results at *Z*_*th*_=0: one-sample t-tests for controls (CN) in A, one-sample t-tests for schizophrenia (SZ) patients in B, and robust regression results showing significant (CN-SZ) group differences in C. Panels D (1-sample CN), E (1-sample SZ), and F (robust regression CN-SZ) show the respective analyses at *Z*_*th*_=0.5. The color bar for one-sample tests (A, B, D, E) indicates the significance and sign of the coupling (k=-log10(p) × sign(t)), with hotter colors for positive coupling and cooler colors for negative coupling. For one-sample tests, only very significant results with (magnitude of k*>*= 10) are shown, while for the group differences(C, F), all significant results are shown. The color bar for group differences (C, F) indicates the significance and direction of group differences.

## 3 Results and Discussion

### 3.1 Dynamic Spatial Brain Networks, Hierarchical Activity, and Complexity

Our analysis identified 14 distinct intrinsic connectivity networks (ICNs), as shown in Figure 2: VIS-P (visual primary), SUB (subcortical), MTR-P (somatomotor primary), CER (cerebellar), ATN (attention-dorsal), FRNT (frontal), MTR-S (somatomotor secondary), FPN-R (frontoparietal right), VIS-S (visual secondary), pDMN (posterior default mode), FPN-L (frontoparietal left), SN (salience), TEMP (temporal), and aDMN (anterior default mode). By visualizing these networks at different activity thresholds, as shown in Figure 2, we can peel away the outer layers to reveal the most active core regions. For example, at a lower threshold (*Z*_*th*_=0), a network like the Primary Visual Network (VIS-P) appears widespread encompassing a large number of voxels with above average activity. How-ever, as we increase the threshold to *Z*_*th*_=2.5, the network visualization shrinks to show only the most highly active, focused core regions. This hierarchical organization of activity within each network is a key aspect of their spatiotemporal dynamics, which our analysis is designed to capture using FrD that can be visualized as a property that varies based on the level of activity examined within the network as illustrated in Figure 3. This shows that networks have a core-periphery structure where more highly active regions exhibit a lower FrD, suggesting a more concentrated, less complex spatial pattern, while the wider network at lower thresholds shows greater complexity. These networks are also spatially dynamic with their complexity changing over time (Figure 3F).

### 3.2 Predominantly higher spatial complexity (mean FrD) and altered spatial focus (mean volume) in SZ

Robust regression results with age, sex and site as covariates comparing the subject-level mean FrD across windows between the CN and SZ groups reveal significant differences across several networks and thresholds, as depicted in Figure 4. The results show that, for most of the networks with significant group differences, the mean FrD is predominantly higher in the SZ group compared to controls (blue) (core of VIS-P, MTR-P, ATN, MTR-S, VIS-S, pDMN, TEMP and periphery of SUB, CER, FPN-R, FPN-L, aDMN). This finding suggests that while CN networks exhibit a more focused, less complex structure, networks in individuals with SZ tend to have more spatially irregular patterns and complex boundaries. On the other hand, the core of SUB, CER, FPN-R, FPN-L and the periphery of pDMN and TEMP show lower complexity in SZ compared to CN. Similar analysis results with mean volume shown in Figure 5 reveal that several networks become less spatially focused (more widespread) in SZ with lower core volumes and higher volumes near the periphery (SUB, CER, FRNT, FPN-R, FPN-L, aDMN) while several others become more spatially focused in SZ with higher core volumes and lower volumes near the periphery (ATN, VIS-S, pDMN, TEMP). Together, these results highlight the altered spatial focus with higher complexity in SZ.

### 3.3 Reduced Dynamical Flexibility (std. of FrD) in SZ

We investigated the temporal variability of the network’s spatial complexity by comparing the standard deviations of FrD and volume of networks across windows between the two groups. As summarized in Figure 6, the results consistently show that the standard deviation of FrD is lower in the SZ group compared to CN across multiple networks and thresholds (VIS-P, SUB, MTR-P, CER, ATN, FRNT, MTR-S, FPN-R, VIS-S, pDMN, FPN-L, SN, TEMP, aDMN). This reduction is even more pronounced when looking at the std. of volume of these networks in Figure 7. The consistent reduction in the variability of FrD/volume in SZ suggests a loss of dynamical flexibility in these networks, indicating that they undergo less changes in their volume or spatial complexity over time.

### 3.4 Fractal Dimension Coupling (FrDC)

Our analysis demonstrated that networks exhibit significant FrDC, indicating that their complexity changes in a synchronized fashion. As detailed in Table 1 and Figure 8A,B,D,E, a large number of network pairs across all subjects showed significant non-zero FrDC in 1-sample t-tests, which was predominantly positive at lower thresholds (when the periphery of networks is included). For example, Figure 8A,D show that the network pairs (MTR-P/MTR-S), (VIS-P/VIS-S), (FPN-R/FPN-L), (SN/SUB) have the strongest positive coupling in CN at *Z*_*th*_=0 and *Z*_*th*_=0.5, with synchronized changes in complexity. A comparison between the CN and SZ groups revealed that while the total number of significant FrDC pairs is similar in Table 1, the specific network pairs with significant coupling differ.

**Table 1:** 1-sample t-tests of subject level fractal dimension coupling (FrDC) of network pairs. Number of network pairs (out of 91 possible combinations among 14 networks) with significant non-zero positive (+ve, red) or negative (-ve, blue) FrDC as found from 1-sample t-tests for all subjects, CN, and SZ groups across different activity thresholds (*Z*_th_). The first value in each cell corresponds to significance with 5% FDR correction at a single threshold (91 comparisons), while the second (bold) value corresponds to significance with 5% FDR correction across all thresholds (a total of 91 × 8 comparisons). FrDC is predominantly positive at lower thresholds (which include network periphery).

| $Z_{th}$ | AllSubjects +ve | AllSubjects -ve | CN +ve | CN -ve | SZ +ve | SZ -ve |
| --- | --- | --- | --- | --- | --- | --- |
| 0.0 | 67, 67 | 4, 3 | 56, 56 | 3, 3 | 67, 64 | 2, 1 |
| 0.5 | 52, 52 | 8, 8 | 46, 46 | 14, 14 | 52, 52 | 1, 1 |
| 1 | 91, 91 | 0, 0 | 91, 91 | 0, 0 | 91, 89 | 0, 0 |
| 1.5 | 55, 54 | 11, 11 | 49, 49 | 13, 13 | 47, 47 | 5, 5 |
| 2 | 36, 36 | 29, 29 | 33, 33 | 28, 27 | 29, 29 | 14, 16 |
| 2.5 | 30, 30 | 30, 32 | 29, 29 | 28, 28 | 22, 22 | 13, 14 |
| 3 | 27, 27 | 28, 28 | 25, 25 | 25, 25 | 18, 20 | 9, 12 |
| 3.5 | 29, 31 | 6, 8 | 23, 26 | 8, 9 | 14, 18 | 1, 2 |

Further investigation with robust regression analysis (Table 2 and Figure 8C,F) showed that several network pairs have significantly different FrDC between the two groups. At *Z*_*th*_=0, we found significant group differences in the FrDC for the pairs (ATN/SUB), (MTR-S/SUB), and (VIS-S/FPN-R). At *Z*_*th*_=0.5, we identified significant differences in the FrDC of the pairs (SUB/VIS-P), (ATN/SUB), and (VIS-S/FPN-R). A key finding, as noted in Table 2 with the corresponding 1-sample t-test results, is that all network pairs showing significant group differences have a higher positive FrDC in the SZ group compared to CN, which has either neutral or negative coupling. This points to an aberrant FrDC pattern in SZ.

**Table 2:** Group differences (CN-SZ) in subject level fractal dimension coupling (FrDC) of network pairs. The table lists network pairs with significant group differences in FrDC between CN and SZ groups, as seen with robust regression with age, sex and site as covariates, using 5% FDR correction at each threshold. All values shown are (k=-log10(p) × sign(t)). The data suggests that all network pairs with significant (CN-SZ) group differences have a higher positive FrDC (blue) in the SZ group compared to the CN group, which has either neutral or negative coupling, as revealed by the corresponding 1-sample t-test results in the table. Significant results at only the lower thresholds imply that these changes are focused near the periphery of these networks.

| $Z_{th}$ | ICN<sub>1</sub> | ICN<sub>2</sub> | Regression results (CN-SZ) | 1-sample t-test (all subjects) | 1-sample t-test (CN) | 1-sample t-test (SZ) |
| --- | --- | --- | --- | --- | --- | --- |
| 0 | ATN | SUB | -2.9121 | 2.1377 | 0.020937 | 4.51 |
|  | MTR-S | SUB | -3.3173 | 3.8699 | 0.36715 | 6.9856 |
|  | VIS-S | FPN-R | -3.6482 | 1.6301 | -0.11873 | 4.2007 |
| 0.5 | SUB | VIS-P | -3.7917 | -0.23687 | -2.4893 | 2.2061 |
|  | ATN | SUB | -2.8363 | 0.71845 | -0.53718 | 2.9011 |
|  | VIS-S | FPN-R | -3.8565 | -0.47811 | -2.9438 | 1.6485 |

### 3.5 Association of FrD Metrics with Subject Scores

Robust linear regression analysis was performed to examine the links between FrD metrics and subject cognitive (visual learning, verbal learning, processing speed and working memory), symptom (positive and negative syndrome scale - PANSS -total symptom score for patients), and chlorpromazine (CPZ) drug scores for patients. The results, as summarized in Table 3, show a strong association of FrD metrics (mean FrD, std FrD, FrDC) with scores, with several networks exhibiting significant positive or negative associations with Visual Learning, Verbal Learning, Processing Speed and PANSS total. Associations with FrDC were found to be very rare. The only significant association was with Working Memory and the (TEMP/aDMN) network pair at *Z*_*th*_=0. Working Memory or CPZ Drug scores did not show any other significant associations.

**Table 3:**
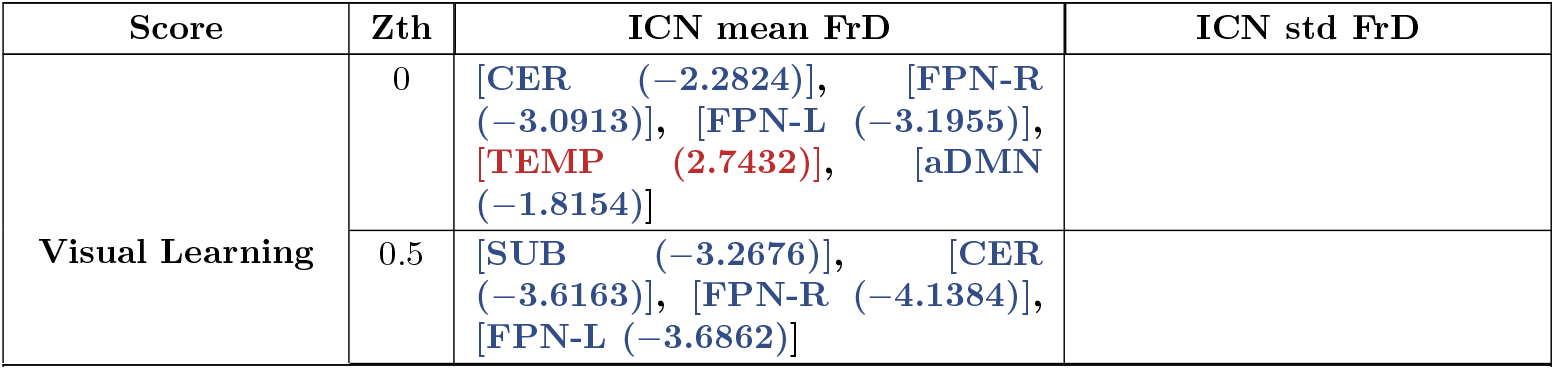
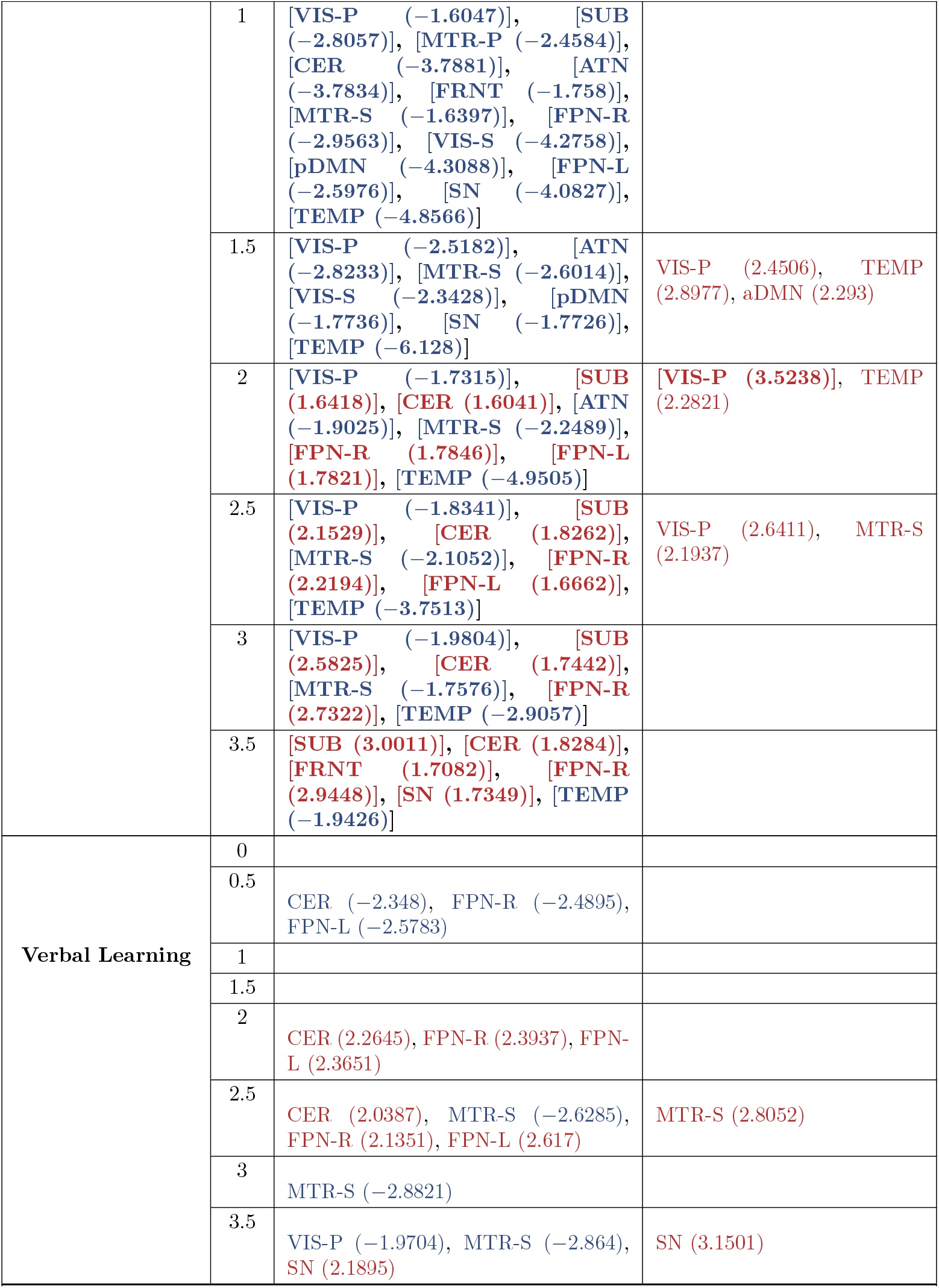
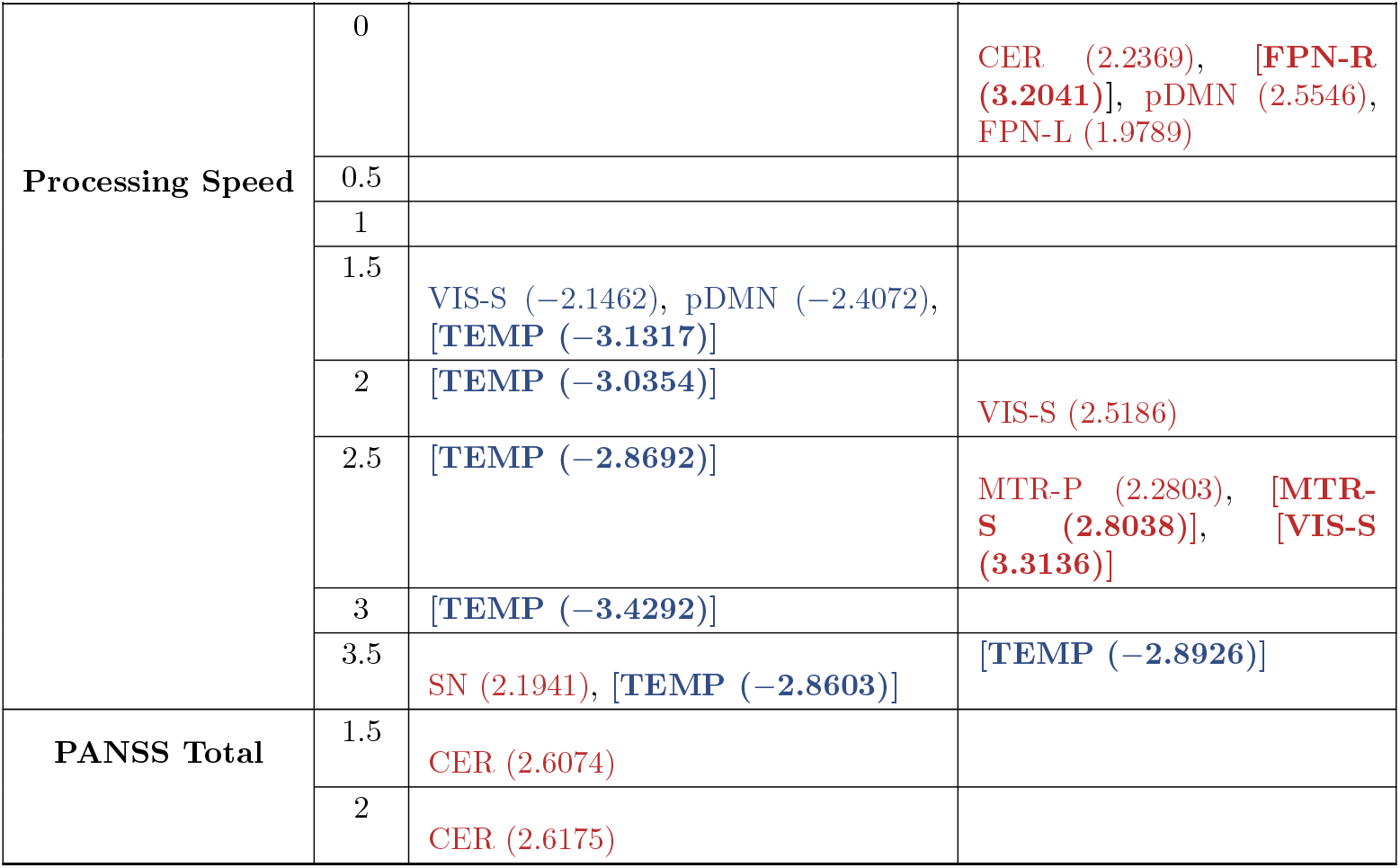
Association of subject cognitive, symptom, drug scores with subject level FrD metrics. The table lists network pairs with significant associations with subject scores. All values shown are (k=-log10(p) × sign(t)), where t is the ratio of the regression coefficient to the standard error of its estimate. Predictors with positive coefficients are shown in red, while those with negative coefficients are shown in blue. Significant results with 5% FDR correction for multiple comparisons at each threshold are shown and results significant with FDR across all thresholds are shown in bold. Associations with FrDC are very rare and only found for Working Memory with (TEMP, aDMN) network pair at *Z*_*th*_ = 0 (not shown here). No significant associations are found for any of the scores with FrDC of any other network pair at any of the thresholds. No other significant associations were found with Working Memory or CPZ Drug scores. The predominant association of all the cognitive scores with std(FrD) is positive, meaning more dynamic flexibility results in higher scores, with std(FrD) noted earlier (Fig.6) to be consistently higher in CN than SZ across networks.

| Score | Zth | ICN mean FrD | ICN std FrD |
| --- | --- | --- | --- |
| Visual Learning | 0 | [CER (-2.2824)], [FPN-R (-3.0913)], [FPN-L (-3.1955)], [TEMP (2.7432)], [aDMN (-1.8154)] |  |
|  | 0.5 | [SUB (-3.2676)], [CER (-3.6163)], [FPN-R (-4.1384)], [FPN-L (-3.6862)] |  |
|  | 1 | [VIS-P (−1.6047)], [SUB (−2.8057)], [MTR-P (−2.4584)], [CER (−3.7881)], [ATN (−3.7834)], [FRNT (−1.758)], [MTR-S (−1.6397)], [FPN-R (−2.9563)], [VIS-S (−4.2758)], [pDMN (−4.3088)], [FPN-L (−2.5976)], [SN (−4.0827)], [TEMP (−4.8566)] |  |
|  | 1.5 | [VIS-P (−2.5182)], [ATN (−2.8233)], [MTR-S (−2.6014)], [VIS-S (−2.3428)], [pDMN (−1.7736)], [SN (−1.7726)], [TEMP (−6.128)] | VIS-P (2.4506), TEMP (2.8977), aDMN (2.293) |
|  | 2 | [VIS-P (−1.7315)], [SUB (1.6418)], [CER (1.6041)], [ATN (−1.9025)], [MTR-S (−2.2489)], [FPN-R (1.7846)], [FPN-L (1.7821)], [TEMP (−4.9505)] | [VIS-P (3.5238)], TEMP (2.2821) |
|  | 2.5 | [VIS-P (−1.8341)], [SUB (2.1529)], [CER (1.8262)], [MTR-S (−2.1052)], [FPN-R (2.2194)], [FPN-L (1.6662)], [TEMP (−3.7513)] | VIS-P (2.6411), MTR-S (2.1937) |
|  | 3 | [VIS-P (−1.9804)], [SUB (2.5825)], [CER (1.7442)], [MTR-S (−1.7576)], [FPN-R (2.7322)], [TEMP (−2.9057)] |  |
|  | 3.5 | [SUB (3.0011)], [CER (1.8284)], [FRNT (1.7082)], [FPN-R (2.9448)], [SN (1.7349)], [TEMP (−1.9426)] |  |
| Verbal Learning | 0 |  |  |
|  | 0.5 | CER (−2.348), FPN-R (−2.4895), FPN-L (−2.5783) |  |
|  | 1 |  |  |
|  | 1.5 |  |  |
|  | 2 | CER (2.2645), FPN-R (2.3937), FPN-L (2.3651) |  |
|  | 2.5 | CER (2.0387), MTR-S (−2.6285), FPN-R (2.1351), FPN-L (2.617) | MTR-S (2.8052) |
|  | 3 | MTR-S (−2.8821) |  |
|  | 3.5 | VIS-P (−1.9704), MTR-S (−2.864), SN (2.1895) | SN (3.1501) |
| Continued on next page |  |  |  |
| Processing Speed | 0 |  | CER (2.2369), [FPN-R (3.2041)], pDMN (2.5546), FPN-L (1.9789) |
|  | 0.5 |  |  |
|  | 1 |  |  |
|  | 1.5 | VIS-S (−2.1462), pDMN (−2.4072), [TEMP (−3.1317)] |  |
|  | 2 | [TEMP (−3.0354)] | VIS-S (2.5186) |
|  | 2.5 | [TEMP (−2.8692)] | MTR-P (2.2803), [MTR-S (2.8038)], [VIS-S (3.3136)] |
|  | 3 | [TEMP (−3.4292)] |  |
|  | 3.5 | SN (2.1941), [TEMP (−2.8603)] | [TEMP (−2.8926)] |
| PANSS Total | 1.5 | CER (2.6074) |  |
|  | 2 | CER (2.6175) |  |

Notably, the predominant association of all cognitive scores with the standard deviation of FrD is positive (red), meaning that greater dynamic flexibility (higher std. FrD) is linked to improved scores. This finding, combined with the observation from Figure 6 and Figure 7 that SZ patients have consistently lower std. FrD (and std. volume), suggests that reduced dynamical flexibility may be a mechanism underlying cognitive impairment in SZ. In addition, at lower thresholds that include the periphery of networks, the mean FrD is predominantly associated negatively (blue) with cognitive scores, implying higher network complexity is related to reduced performance. Combined with the results in Figure 4, where SZ shows higher network complexity (blue), we may associate the increased complexity in these networks to reduced cognitive performance in SZ.

## 4 Discussion

Our comprehensive analysis, utilizing both volumetric measures and fractal dimension (FrD), provides novel insights into the disruption of intrinsic spatial properties in schizophrenia (SZ). While volume quantifies the network’s spatial extent (its capacity for expansion and contraction), FrD provides a measure of its intrinsic spatial complexity and structural organization. The finding that the mean FrD is predominantly higher in SZ patients suggests more spatially irregular patterns and complex network structure compared to the more focused, less complex network patterns found in healthy controls. This is accompanied by altered spatial focus of activity as seen in mean volume changes. This aligns with the idea of a less efficient or a more disorganized network architecture in SZ, where activity may be less confined to specific functional regions and more spatially irregular. This finding is consistent with and extends previous studies that have shown altered network modularity and reduced efficiency in SZ [22, 23, 24]. This observed disruption in spatial complexity can be viewed through the lens of a core-periphery model of brain networks. In healthy brains, networks are thought to have a stable, highly connected core that performs specialized functions and a more flexible periphery that can adapt to changing demands. Our findings could suggest a breakdown of this core-periphery structure, with the network structure becoming less focused with more irregular patterns. This could disrupt the network’s ability to efficiently process and transmit information.

Perhaps more critically, the highly consistent finding of reduced standard deviation in both volume and FrD across all intrinsic connectivity networks at several thresholds in the SZ group suggests a synergistic loss of dynamic flexibility in these networks, resulting in spatial rigidity in the SZ brain. We refer to this pathological convergence as maladaptive complexity. The networks are not only ‘stuck’ in a rigid state struggling to change their spatial extent and complexity to meet the demands of different cognitive states, as evidenced by the low variability of FrD/volume, but they are also ‘stuck’ in an inherently aberrant or disorganized configuration, as evidenced by the predominantly higher mean FrD and altered mean volume. This combined finding may represent a key measurable mechanism for the cognitive deficits and disorganized thought seen in schizophrenia.

Our regression analysis supports this, showing that a higher standard deviation of FrD is associated with improved cognitive scores and a higher mean FrD in network periphery is associated with a drop in scores. These results combined with similar associations between volume metrics and cognitive scores found in our previous study [7] suggest that the brain’s ability to fluidly change its network structure and complexity is crucial for cognitive performance. The observed reduction in this variability in SZ and the breakdown of core-periphery structure could therefore be a key mechanism underlying the cognitive deficits associated with the disorder. This finding resonates with the concept of brain criticality, which posits that a healthy brain operates at a delicate balance between highly ordered and highly random activity [25, 26]. The reduced variability of FrD/volume in SZ could indicate that the brain is “stuck” in a less-than-optimal state, unable to fluidly reorganize itself, a finding also supported by studies showing decreased complexity of brain signals in SZ [27].

Our findings of altered FrD and reduced dynamic flexibility in SZ are consistent with, and extend, previous research on reduced functional connectivity and network dysregulation in the disorder [2, 4]. The dysfunction of specific networks, such as the visual networks, aligns with a body of research demonstrating visual processing deficits in SZ [28, 29]. The reduced dynamic flexibility of these networks might hinder the brain’s ability to fluidly process and integrate complex visual information, which could contribute to visual disturbances and even hallucinations. Similarly, the altered complexity and coupling in the default mode network (DMN) and fronto-parietal networks (FPNs) suggest a breakdown in their fundamental dynamic interplay. An inability to decrease DMN complexity or variability when an FPN needs to take over could lead to the intrusive self-referential thoughts and cognitive disorganization often seen in SZ [30]. Our findings of altered FrDC between networks such as VIS-S/FPN-R and ATN/SUB further supports the dysconnectivity hypothesis [31], demonstrating that disruptions in brain-wide networks and their coupling are associated with the psychopathology and altered behaviors seen in SZ [32]. The increased positive FrDC between networks like SUB, CER and others in SZ could reflect an overly synchronized and potentially less efficient pattern of activity. While a healthy brain’s networks operate with a degree of independence, this increased coupling in SZ might represent a loss of functional segregation, where distinct networks lose their specialized roles and become “stuck” in a single, rigid state.

Our work on FrD can be positioned as a complementary approach to traditional graph theory, which typically describes the relationships between nodes. By characterizing the intrinsic spatial organization and complexity of the networks themselves, our study adds another layer of analysis to the “dysconnectivity” hypothesis. We find that the brain’s ability to be flexible in its complexity, or its “dynamic adaptability,” is disrupted in SZ, and this may provide a measurable mechanism for a key clinical feature of the disease: the inability to respond to changing environments. Our fractal complexity approach measures the quality of a network’s organization and can provide a quantitative explanation for how some patients with SZ struggle with a wide range of tasks and situations. This could manifest as overuse of random exploration strategies, a lack of directed, goal-oriented behavior. This may also be linked to disrupted cognitive processes and reduced sensitivity to rewards, where the brain’s failure to generate diverse and flexible patterns of activity leads to fixed, rigid thought processes and behaviors that are characteristic of the disorder.

## 5 Conclusions

In this study, we investigated the fractal dimension of dynamic spatial brain networks and their coupling to characterize network complexity in schizophrenia. Our results reveal that schizophrenia is associated with several key alterations in their properties:

- **Maladaptive complexity:** Networks in schizophrenia patients tend to show more spatially irregular and complex patterns than those in healthy controls. Schizophrenia is also characterized by a significant reduction in the temporal variability or dynamic flexibility of network complexity.
- **Aberrant connectivity:** The pattern of synchronized complexity changes between networks as seen with FrDC is aberrant in schizophrenia, with patients showing a higher positive FrDC for several network pairs.
- **Association with cognition:** Dynamic spatial brain network complexity features show strong association with cognitive scores with maladaptive complexity in schizophrenia offering a potential mechanism for cognitive deficits.

Overall, our findings highlight the importance of studying the overlooked spatial dynamics of brain networks. The reduced dynamic flexibility of these networks appears to be a robust marker for schizophrenia and may contribute to the cognitive impairments seen in the disorder. Our dSNCC framework could provide a novel, quantifiable lens for immediate application to understand other complex disorders characterized by network dysfunction, such as Alzheimer’s disease, autism, or major depressive disorder. Future work should further explore the interplay between functional and spatial dynamics to provide a more comprehensive understanding of the brain’s complex and ever-changing landscape. For instance, future research could directly test the hypothesis of reduced dynamic adaptability by comparing network FrD/volume at rest with FrD/volume during a cognitive task. This would provide a direct measure of the brain’s ability to change its complexity and spatial properties in response to a specific challenge. Furthermore, the relationship between FrD and specific symptoms like disorganized thought or anhedonia could be explored to find targeted biomarkers. A critical next step will be to bridge our macro-level findings (dSNCC metrics) with micro-level pathology, integrating multi-omics data (e.g., transcriptomics and genetics) or specific molecular imaging to test if the observed network rigidity is mechanistically driven by specific cell-type vulnerability or known glutamatergic or dopaminergic system dysfunctions implicated in schizophrenia and other mental disorders [33, 34].

## 6 Acknowledgements

This research was partially supported by the NSF (grant 2112455) and the NIH (grants R01MH123610 to Dr. Vince D. Calhoun and 5R01MH119251 to Dr. Armin Iraji). We are grateful to the members of the TReNDS Center for their valuable discussions.

## 7 Author Contributions

K.P. and V.D.C. conceived the study. K.P., A.I., and V.D.C. designed the dSNCC methodology and analyses. A.B.A. provided clinical consultation and insight. K.P. performed the data analysis and prepared the figures. K.P. and V.D.C. wrote the main manuscript text. A.I. and A.B.A. reviewed and edited the manuscript. All authors reviewed the results and approved the final manuscript.

